# Induced pluripotent stem cell-derived macrophages as a model for human inflammasome signaling

**DOI:** 10.1101/2025.09.05.674456

**Authors:** Chloe M. McKee, Melanie Cranston, Emma C. McKay, Mohammad Arefian, Thea J. Mawhinney, Ben C. Collins, Rebecca C. Coll

## Abstract

Macrophage models are a mainstay of inflammasome research, however current human *in vitro* macrophage models have significant limitations. Here we generate induced pluripotent stem cell (iPSC)-derived macrophages (iMacs) to study inflammasome signaling and benchmark them with human monocyte-derived macrophages (HMDMs). We confirm that iMacs express high levels of macrophage markers and are highly phagocytic. Whole cell proteomics analysis shows that iMacs express many inflammasome sensors and related proteins, and in functional assays iMacs respond to multiple inflammasome stimuli. The NLRP3 inflammasome is strongly activated in iMacs and we find that nigericin alone activates NLRP3. The non-canonical inflammasome does not require a priming step in iMacs as caspase-4 is constitutively expressed. High levels of NAIP/NLRC4 inflammasome activation are also observed in response to needle toxin. Finally, unlike HMDMs, iMacs activate NLRP1. Therefore, we demonstrate that iMacs are a physiologically relevant and highly tractable model to study human inflammasome signaling and regulation.

**Motivation:** iPSC-derived macrophages (iMacs) are functionally, transcriptionally, and phenotypically similar to primary human macrophages. iMacs therefore offer new opportunities to study inflammasome activity in a human macrophage model, but to date they have not been widely used. In this study, we describe a protocol to differentiate and characterize iMacs. We then describe how to activate a range of different inflammasomes within these cells and assess the inflammasome response by measuring pyroptosis, cytokine release, ASC speck formation, and processing of inflammasome-related proteins. We also benchmark iMac responses with the current gold standard primary human monocyte derived macrophage model.

## Introduction

Macrophages are a key component of the innate immune system that express a wide range of pattern recognition receptors (PRRs). Cytosolic PRRs such as nucleotide oligomerization domain (NOD)-like receptors (NLRs) can assemble into multi-protein complexes termed ‘inflammasomes’(1). Inflammasomes are activated by a range of microbial-, host-, and environmental-derived stimuli to trigger a potent inflammatory response(2). Inflammasome activation induces recruitment of the adapter protein apoptosis associated speck-like protein containing a CARD (ASC). ASC filaments provide a platform to activate the effector protein caspase-1 which cleaves the pro-inflammatory cytokines interleukin (IL)-1β and IL-18 and causes gasdermin D (GSDMD)-mediated inflammatory cell death (pyroptosis)(2). Aberrant activation of this highly inflammatory signaling program is associated with a range of inflammatory diseases and inflammasome inhibitors are of significant therapeutic interest. Indeed, several NOD-, LRR-, and pyrin domain-containing protein 3 (NLRP3) inflammasome inhibitors are in clinical trials for conditions including cryopyrin associated periodic syndromes (CAPS), asthma, and Parkinson’s disease(3,4). Inflammasome receptors, such as NLRP3 and NLR family CARD domain-containing protein 4 (NLRC4) are expressed primarily by macrophages and other myeloid cells(5). Therefore, macrophages are the main cell model used in the study of inflammasome function. However, current *in vitro* macrophage models have significant limitations.

Primary mouse bone marrow-derived macrophages (BMDMs) are widely used, but significant differences exist in immune development and the innate and adaptive immune systems between mice and humans(6). For example, mice are less sensitive to lipopolysaccharide (LPS) than humans and, although the regulation of many LPS-responsive genes is conserved between mice and humans, 24% of orthologs are divergently regulated(7,8). Importantly, there are significant differences in the expression, structure, and activation of inflammasomes in humans and mice. Up to seven NLRP1 paralogs have been described in different mouse strains while humans express a single NLRP1 gene and a related inflammasome sensor, CARD8, does not have a mouse homolog(9,10). Mice express seven NAIPs that trigger activation of NLRC4, while humans express a single functional NAIP that recognizes bacterial type III secretion system (T3SS) proteins(11–15). In addition, regulatory proteins such as caspase recruitment domain (CARD)-only and pyrin domain (PYD)-only proteins (COPs and POPs) are not expressed in mice, therefore, COPs and POPs perform functions in fine tuning primate immune responses that are absent in rodents(16). Interestingly, we recently identified small molecule NLRP3 inhibitors, BAL-0028 and BAL-0598, that are highly effective in human and primate cells but have significantly reduced activity in mouse cells(17). To explore the potential clinical benefits of targeting inflammasomes, it is critical to understand inflammasome regulation in human systems.

Current human models include the human monocytic leukemia cell line THP-1 which can be differentiated into a macrophage-like state using phorbol-12-myristate-13-acetate (PMA)(18). However, THP-1s are functionally immature, have karyotypic abnormalities, and the expression of many cytokines and chemokines, such as IL-8, interferon (IFN)-γ, and monocyte chemoattractant protein-1, are dysregulated in THP-1s(19,20). Importantly, PMA differentiation increases pro-IL-1β and NLRP3 expression, confounding interpretation of inflammasome assays(21). Primary human monocyte-derived macrophages (HMDMs) are considered the gold standard human macrophage model(22) . However, there are practical and ethical issues regarding the frequency and volume of blood donations that can be taken as HMDMs are non-proliferative(19,22). Additionally, the monocyte isolation techniques used to generate HMDMs can impact their polarization, phenotype, purity, and cell yield(23). Experiments with HMDMs are additionally challenging as responses can vary significantly due to the myriad inherent differences between donors such as age, sex, and genetic heterogeneity. HMDMs are also difficult to genetically modify as they mount a robust immune response against foreign nucleic acids(24). Finally, HMDMs are not truly representative of tissue-resident macrophages as they have different embryological origins(25). Therefore, new *in vitro* human macrophage models overcoming these limitations would be useful for studying inflammasomes.

Human iPSCs have emerged as a powerful tool to model human development and disease across many cell types including immune cells like macrophages. Since the first study for differentiating iPSCs into macrophages (iMacs) was published in 2011, various protocols have been developed, including co-culture with mouse stromal fibroblasts or embryoid body (EB) formation through mechanical dissociation(26–29). More recent protocols, such as Douthwaite et al (2022), induce EB formation by centrifugation of low-adherent 96 well plates with bone morphogenetic protein 4 (BMP-4), then induce hematopoiesis with IL-3 and macrophage colony-stimulating factor (M-CSF) to generate myeloid precursor cells, which are further differentiated into macrophages with M-CSF(30). iMacs generated with these recent differentiation protocols express a wide range of macrophage specific markers, including CD14, CD16, CD44, CD45, CD64, CD68, CD195, and CD200(22,31). iMacs also display functional activity with conserved responses to inflammatory stimuli and high levels of phagocytosis and antimicrobial activity(24,31). These protocols offer a continuous and scalable supply of cells that can be genotype- and patient-specific(22,29,32). iPSCs are very amenable to genetic manipulation as they are immunologically unresponsive and can therefore be genetically modified before differentiation into macrophages possessing the same mutation(22). iMacs are also an attractive alternative to BMDMs as the use of insentient, *in vitro* material adheres to the 3Rs concept to replace the use of living animals(33) . iMacs therefore offer new opportunities in studying macrophage biology, including inflammasome regulation.

To date, few published studies look at inflammasome responses in iMacs. Staphylococcal pore-forming toxins have been shown to activate NLRP3 in iMacs(34). Also, iMacs differentiated from iPSCs derived from a patient with neonatal onset multisystem inflammatory disease (NOMID) display constitutively activated NLRP3 activity(35). Most recently, iMacs have been shown to activate NLRP3 independently of NIMA Related Kinase 7 (NEK7), with IκB kinase β (IKKβ) instead priming the inflammasome by triggering NLRP3 recruitment to the trans-Golgi network(36). However, there has been no in-depth characterization of other inflammasome responses within iMacs. Inflammasomes play important roles in human disease and infection, with NLRP1 activation triggered in response to viral infection and the NAIP/NLRC4 and non-canonical inflammasomes in response to bacterial infection(11,37,38)(11,37,38). Further investigation was therefore required to determine whether iMacs are useful as a model that is comparable to HMDMs for inflammasome studies.

## Results

### Expression of macrophage markers in iMacs

To generate iMacs, we adapted the Douthwaite et al., (2022) protocol using KOLF2-C1 iPSCs, a skin-derived iPSC line generated from a healthy male of European ancestry by the Sanger institute(30). KOLF2-C1 has been identified as a well-performing iPSC line that was originally reprogrammed using non-integrating methods and in media and substrates that were chemically defined(39). KOLF2-C1 are also widely used in the stem cell field, and they maintain genomic integrity during multiple rounds of gene editing with CRISPR/Cas9(39).

To generate EBs, iPSCs are seeded on low-adherent U-bottomed 96 well plates and stimulated with BMP-4. The plates are centrifuged and incubated for 5 days. The EBs are then transferred to flasks and M-CSF and IL-3 are used to drive the cells towards a myeloid lineage. Within 4-5 days, the EBs adhere to the surface of the flasks and produce a layer of stromal cells, termed the ‘stromal skirt’. After 3 – 4 weeks of culture, the EBs begin to produce myeloid precursors. The myeloid precursors are differentiated into macrophages for 7 days with M-CSF. This protocol is summarized in Supp. Fig 1A.

To confirm that our protocol generates macrophage-like cells, we examined a panel of known macrophage markers by flow cytometry (CD14, CD16, CD44, CD64, and CD68)(30). While monocytes also express most of these markers, CD16 expression is low in monocytes (∼10%)(30). CD14 and CD44 are highly expressed in iMacs (Fig. 1A, B) at levels comparable to HMDMs (Supp. Fig. 1B, Fig. 1C). CD64 and CD68 are also highly expressed in iMacs and HMDMs, although the expression level varied between iMac batches and HMDM donors (Fig. 1B, C). CD16 expression in iMacs is lower than the other markers (Fig. 1A, B) in line with findings from van Wilgenburg et al., (2013) and Douthwaite et al., (2022) who demonstrated that CD16 staining is intermediate in iMacs(29,30). In contrast, HMDMs express higher levels of CD16 (Fig. 1C, Supp. Fig 1B). Therefore, iMacs express a range of macrophage markers, indicating they are macrophage-like.

**Fig. 1.**
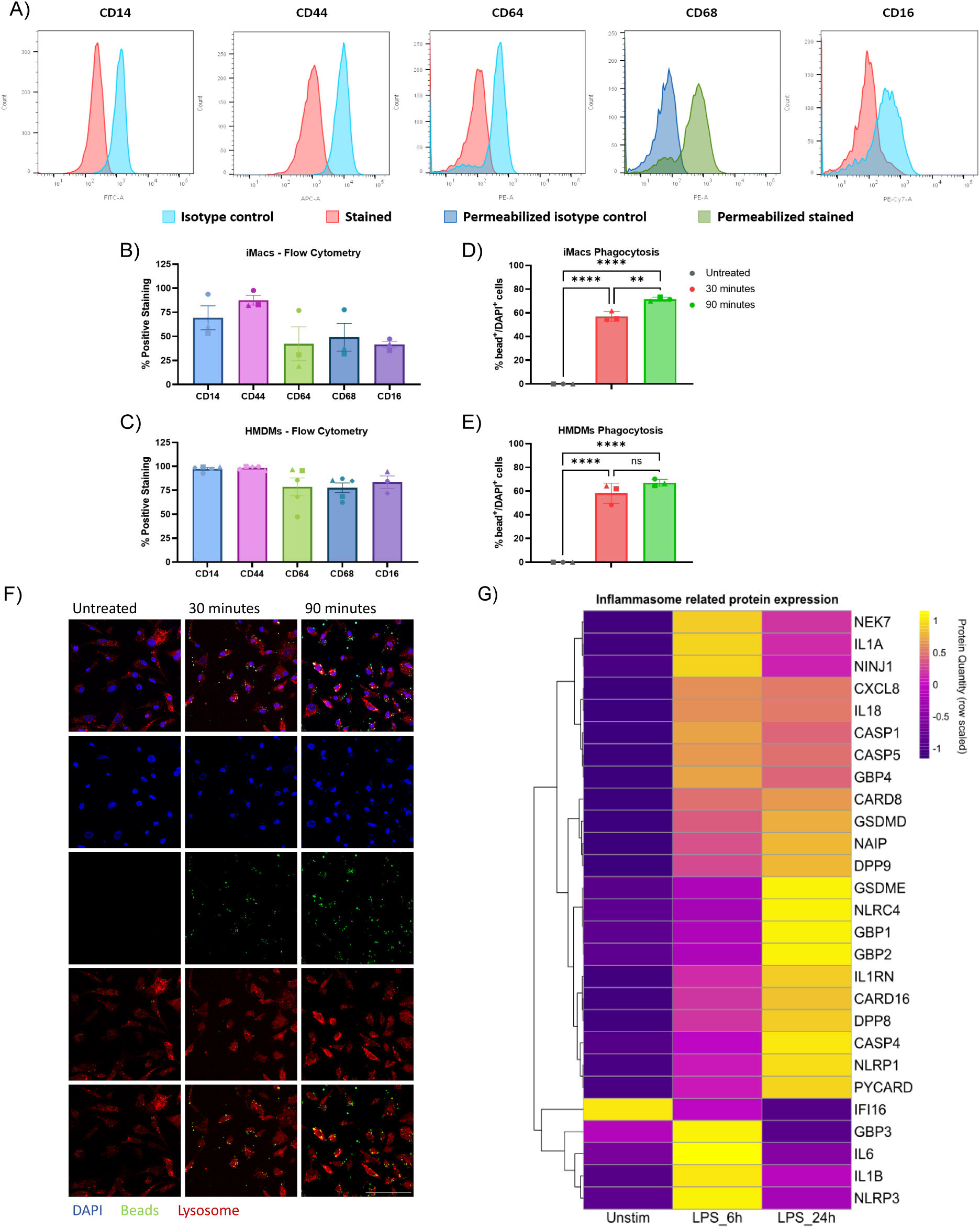
iMacs are macrophage-like with high expression of macrophage markers, phagocytosis and expression of inflammasome components. A) Histograms showing representative expression of macrophage markers CD14, CD44, CD64, CD68, and CD16 in unstimulated iMacs compared to relative isotype controls. Quantification of CD14, CD44, CD64, CD68, and CD16 expression across 3 – 5 independent replicates in B) iMacs and C) HMDMs. Quantification of cells containing at least one fluorescent bead in D) iMacs and E) HMDMs. F) 63 X (oil lens) Stellaris-5 confocal microscope representative images from iMacs in untreated cells and following incubation with Latex amine modified polystyrene fluorescent beads (green) for 30 minutes or 90 minutes stained with DAPI (blue) and Lysotracker (red) with phagocytosed beads highlighted with white boxes. Scale bar = 100 μM. Data are represented as mean +/-SEM, n = 3 independent replicates. **p < 0.01, ****p < 0.0001 by One-way ANOVA with Tukey’s multiple comparisons test. G) Heatmap illustrating scaled protein quantity of inflammasome and inflammation related proteins (rows) following whole cell proteomics analysis of iMacs that were unstimulated or following stimulation with LPS (100 ng/ml) for 6 h or 24 h. Clustering was performed using Euclidean distance and protein quantities are scaled by row.

### Phagocytic activity of iMacs

Macrophages play a vital role in innate immunity by phagocytosing microbes, releasing pro-inflammatory cytokines and antimicrobial factors, and presenting antigens to activate the adaptive immune response(40). To characterize their functionality, we assessed the phagocytic activity of our iMacs. iMacs were treated with latex fluorescent beads for 30 or 90 minutes and lysosomes were stained using LysoTracker-Red (Fig 1 D, F). iMacs are highly phagocytic, with ∼55% of cells containing at least one fluorescent bead after 30 min. This increases to ∼70% of cells after 90 min (Fig. 1D, F). This level of phagocytosis is consistent with HMDMs, where ∼60% cells contain at least one fluorescent bead at 30 minutes and ∼70% after 90 minutes (Fig. 1E, Supp Fig 1C).

### Expression of inflammasome-related proteins

The expression of specific inflammasome sensors varies between cell types, for example airway epithelial cells are believed to express low levels of NLRP3 but high levels of NLRP1(41). Previous studies have not examined global protein expression in iMacs so we performed whole cell proteomic analysis on unstimulated iMacs or following stimulation with LPS for 6 h or 24 h (Supp. Table 1). We observe that iMacs express many inflammasome sensors such as NLRP3, NLRP1, CARD8, NAIP, NLRC4, and caspase-4/5 alongside other inflammasome components such as ASC, caspase-1, GSDMD, NINJ1, and NEK7 (Fig. 1G, Supp. Fig. 2). In agreement with previous studies on NLRP3 priming, NLRP3 expression is enhanced after 6 h LPS but decreased after 24 h LPS(42). The expression of NLRC4 and caspase-4 also increases with LPS stimulation and peaks at 24 h. The iMacs also respond to LPS stimulation by increasing the expression of pro-IL-1β and other inflammasome-independent cytokines such as IL-6 which are upregulated at 6 h LPS. These data demonstrate that iMacs express numerous inflammasome sensors and all the components of the inflammasome pathway.

### Responses to NLRP3 inflammasome stimuli

To functionally characterize iMacs as a model for human inflammasome signaling, we first investigated NLRP3 activation. NLRP3 detects many microbial- and host-derived signals and in macrophages, NLRP3 activation requires an initial priming stimulus followed by a second activation signal(43). To activate NLRP3, iMacs were primed with the bacterial-derived stimuli LPS or Pam3CSK4 which signal through Toll like receptor 4 (TLR4) and TLR1/2 respectively for 4 h. Cells were then treated with the bacterial toxin nigericin or the danger signal extracellular ATP for 2 hours. Inflammasome activity was measured by assessing pyroptotic cell death via lactate dehydrogenase (LDH) release assay, ELISAs for IL-1β and IL-18 release, and western blotting for NLRP3 and pro-IL-1β expression and the cleavage of caspase-1, IL-1β, and GSDMD. As expected, treatment with nigericin triggers pyroptosis in primed iMacs, alongside high levels of active IL-1β release and lower but significant release of IL-18 (Figure 2A & B, Supp. Fig. 3A). In response to LPS and Pam3CSK4, there is also significant release of TNF (Fig. 2C), an inflammasome-independent cytokine induced in response to TLR stimulation. LPS and Pam3CSK4 priming also enhances pro-IL-1β and NLRP3 expression (Fig. 2D). In comparison in HMDMs, LPS or Pam3CSK4 plus nigericin triggers NLRP3 activation but to a lower level than the iMacs, with little cell death and lower levels of IL-1β release, whereas TNF release is comparable between the cell types (Fig. 2E – G). Interestingly, significant cell death is seen in unprimed iMacs treated with nigericin (Fig. 2A), but in HMDMs nigericin alone does not trigger pyroptosis (Fig. 2E). The cleavage of caspase-1 and GSDMD in iMacs stimulated with nigericin alone is similar to primed cells (Fig. 2D), indicating that this is inflammasome-dependent pyroptosis. Treatment with the NLRP3-specific inhibitor MCC950 also blocks nigericin alone-induced pyroptosis (Supp. Fig. 3B), confirming nigericin alone activates NLRP3 in iMacs.

**Fig. 2.**
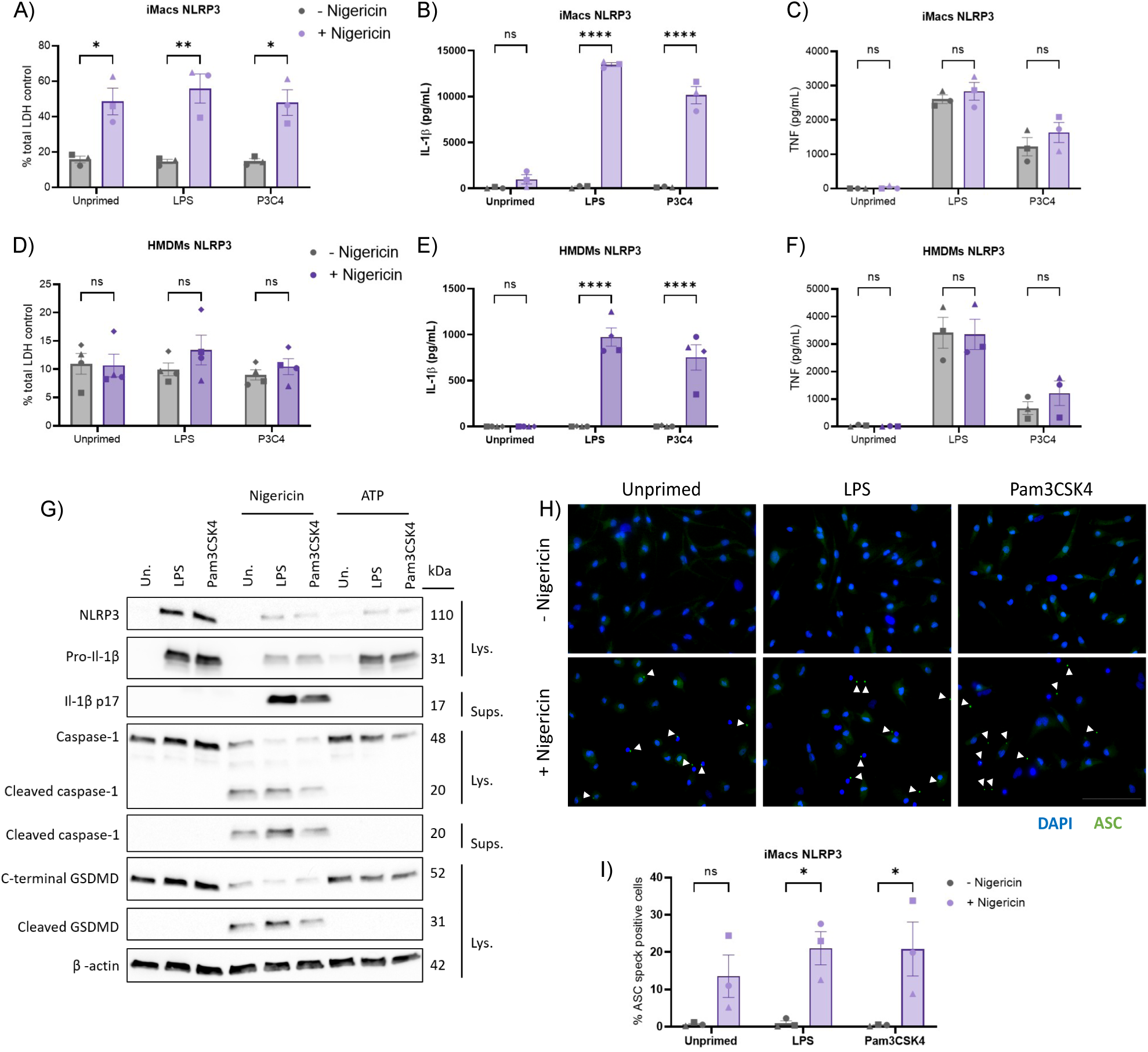
iMacs display higher NLRP3 inflammasome responses compared with HMDMs. Quantification of **A)** LDH release, **C)** IL-1β, and **D)** TNF release, and **D)** Western blots from iMacs in unprimed cells and following stimulation with 100 ng/ml LPS or 1 μg/ml Pam3CSK4 for 4 h with 5 µM nigericin for 2 h. Quantification of **E)** LDH release, F) IL-1β, and **G)** TNF release from HMDMs in unprimed cells and following stimulation with 100 ng/ml LPS or 1 μg/ml Pam3CSK4 for 4 h with 5 µM nigericin for 2 h. **H)** Representative images from iMacs in unprimed cells and following stimulation with 100 ng/ml LPS or 1 μg/ml Pam3CSK4 for 4 h with 5 µM nigericin for 2 h stained with DAPI (blue) and ASC (green) with ASC specks highlighted with white arrows. Scale bar = 100 μM. **I)** Quantification of ASC speck positive cells. Data are represented as mean +/-SEM, n = 3-5 independent replicates, *p < 0.05, **p < 0.01, ***p < 0.001, ****p < 0.0001 by Two-way ANOVA with Šídák’s multiple comparisons test

The response to ATP in iMacs is much lower than nigericin stimulation, with no increase in pyroptosis and lower levels of IL-1β release (Supp. Fig. 3C – E). ATP is equally capable of triggering IL-1β release in HMDMs without pyroptosis, comparable with the iMacs (Supp. Fig. 3F – H).

To confirm formation of the NLRP3 inflammasome in iMacs, ASC specks were visualized by immunofluorescence microscopy. ASC specks, shown in green and highlighted with white arrows, are often located close to the blue 4ʹ,6-diamidino-2-phenylindole (DAPI)-stained nuclei with a single speck per cell (Fig. 2H). Priming followed by nigericin stimulation triggers a significant increase in the percentage of DAPI+ASC+ iMacs (Fig. 2H, I). In unprimed cells treated with nigericin, there is a non-significant increase in the percentage of DAPI+ASC+ cells. The ASC speck formation further confirms that nigericin alone can induce NLRP3 inflammasome formation in iMacs. Together, these data demonstrate that the NLRP3 inflammasome is functional in iMacs and responsive to multiple priming and activation stimuli.

### Response to non-canonical inflammasome stimulation

We next examined non-canonical inflammasome (NCI) activation. In human cells the NCI is activated by cytosolic LPS which triggers caspase-4 and caspase-5 activation and thereby GSDMD-mediated pyroptosis and IL-18 release(37). GSDMD pores also indirectly activate NLRP3 via potassium efflux, thus activating caspase-1 and IL-1β release(44). To study NCI activation, iMacs were primed overnight with Pam3CSK4 before transfection with LPS for 5 h. A control with extracellular LPS alone is included to confirm that responses are due to the transfection of LPS into the cytoplasm. LPS transfection causes increased pyroptosis, IL-1β release, and cleavage of caspase-1, caspase-4, and GSDMD in unprimed iMacs (Fig. 3A-C). These responses are blocked by the caspase-1/4 inhibitor VX-765. NLRP3 inhibition with MCC950 partially reduces IL-1β release and caspase-1 cleavage in LPS transfected cells but does not affect cell death or caspase-4 cleavage, consistent with previous findings in HMDMs by Ross et al., (2018)(44).

**Fig. 3.**
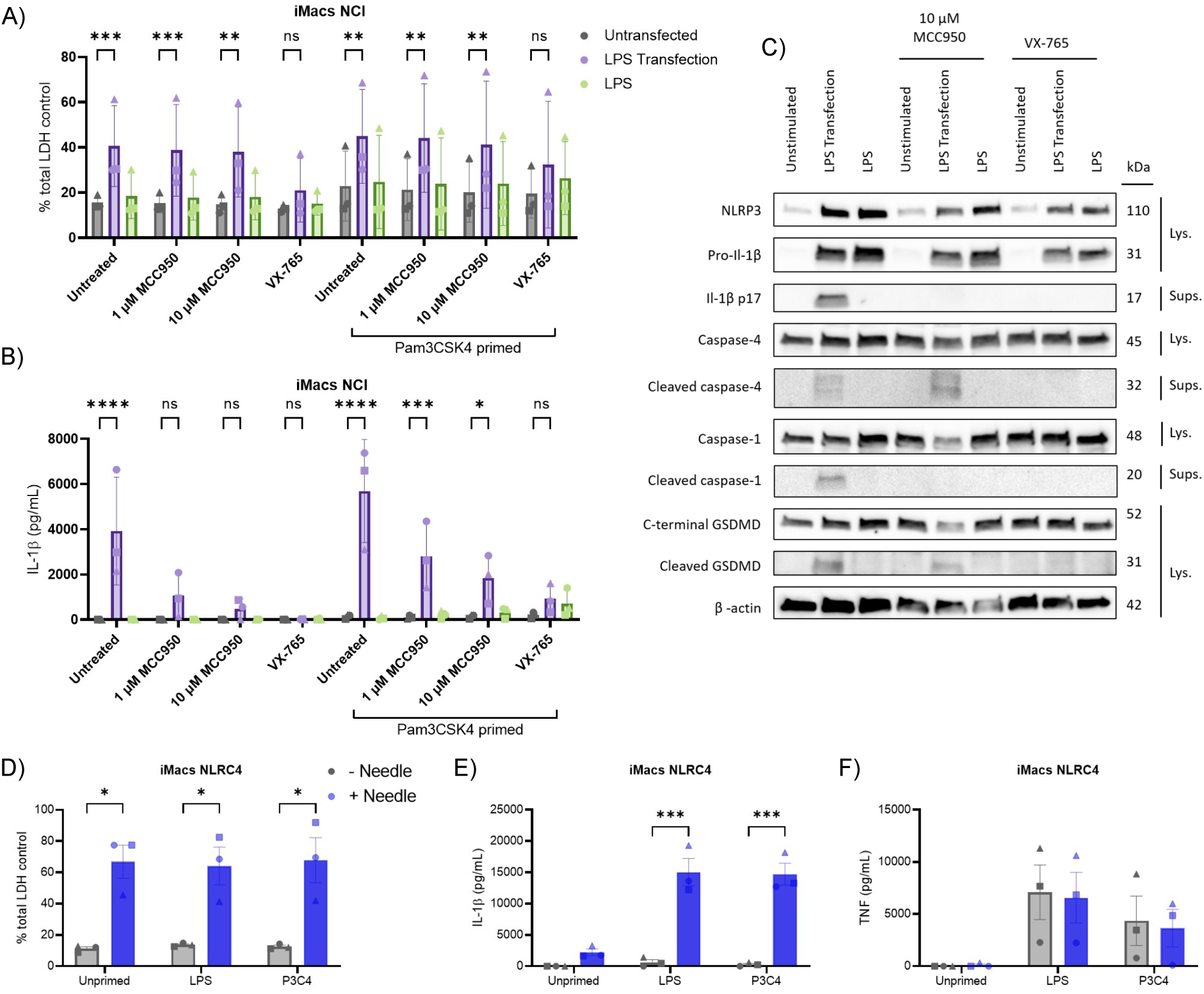
NCI and NLRC4 inflammasome activation in iMacs. Quantification of **A)** LDH and **B)** IL-1β release from iMacs in unprimed and Pam3CSK4 primed cells (1 μg/ml, 16 hours) and **C)** Western blots in from unprimed iMacs following LPS transfection or LPS stimulation for 5 hours and treated with MCC950 (1 μM or 10 μM) or VX-765 (20 μM) for 30 min prior to transfection. Quantification of **D)** LDH, **E)** IL-1β, and **F)** TNF release from iMacs in unprimed cells and following stimulation with 100 ng/ml LPS or 1 μg/ml Pam3CSK4 for 4 h with Lfn-BsaL (1 ng/ml) plus anthrax protective antigen (PA, 250 ng/ml) for 2 hours. Data are represented as mean +/-SEM, n = 3 independent replicates, *p < 0.05, ***p < 0.001, ****p < 0.0001 by repeated measures ANOVA with Tukey’s multiple comparisons test.

All responses are similar in Pam3CSK4 primed iMacs but there is increased IL-1β release due to enhanced pro-IL-1β levels (Fig. 3 A, B, Supp. Fig. 4A). In unprimed iMacs, increased pro-IL-1β and NLRP3 expression is observed both in cells transfected with LPS or treated with extracellular LPS (Fig. 3C). This may explain why IL-1β release is seen in unprimed cells as transfected LPS can also prime iMacs within the 5 hour transfection time.

The response to NCI activation in HMDMs is similar to iMacs as LPS transfection triggered increased cell death and IL-1β release which is inhibited with VX-765 (Supp. Fig. 4B, C). However, priming with Pam3CSK4 appears to be more necessary in HMDMs as IL-1β release in unprimed LPS-transfected cells is lower than in iMacs, indicating iMacs are more sensitive to LPS.

### Response to NAIP/NLRC4 inflammasome stimulation

In humans, NAIP senses bacterial invasion through the detection of components of the T3SS such as needle, inner rod protein, and flagellin. NAIP then associates with NLRC4 to trigger NAIP/NLRC4 inflammasome formation(45). For NAIP/NLRC4 inflammasome activation, iMacs were primed with LPS or Pam3CSK4 and then treated with Lfn-BsaL, a fusion protein of the N-terminal domain of the Bacillus anthracis lethal factor and BsaL needle protein from *Burkholderia pseudomallei*, plus anthrax protective antigen (PA). Lfn-BsaL plus PA alone is sufficient to induce significant pyroptosis in iMacs (Fig. 3D). Priming does not enhance Lfn-BsaL-induced cell death, but in primed cells Lfn-BsaL induces significant IL-1β release (Fig. 3E). TNF release is also induced in response to LPS and Pam3CSK4 (Fig. 3F). The responses in iMacs are very similar to those observed in matched experiments with HMDMs. Similar levels of cell death are observed in response to Lfn-Bsal in HMDMs and they also release significant levels of IL-1β, although IL-1β release is lower than in the iMacs (Supp. Fig. 4D – F). Therefore, NLRC4 activation is similar between iMacs and HMDMs, although iMacs release significantly higher levels of IL-1β.

### Response to NLRP1 and CARD8 inflammasome stimulation

The NLRP1 and CARD8 inflammasomes contain a C-terminal function-to-find domain (FIIND) that must undergo proteolysis, termed ‘functional degradation’, to allow inflammasome activation(46). This can be triggered *in vitro* using inhibitors of the serine proteases dipeptidyl peptidase 8/9 (DPP8/9)(38). To study the response of iMacs to NLRP1/CARD8 stimulation, cells were primed with LPS or Pam3CSK4 and then treated with the DPP8/9 inhibitor Talabostat (Val-boroPro) for 24 hours.

Similar to previous work in T cells(38), priming was not necessary for NLRP1/CARD8 inflammasome activation, as Talabostat alone induces significant pyroptosis (Fig. 4A). There is no significant pyroptosis observed in primed cells treated with Talabostat but there is significant release of IL-1β (Fig. 4B). Priming also triggers significant TNF release (Fig. 4C). These findings in iMacs contrast with those in HMDMs where Talabostat did not induce LDH or IL-1β release (Fig. 4D, E). This is consistent with previous findings(47) and confirms HMDMs are not responsive to NLRP1 activation by Talabostat. The TNF response to LPS and Pam3CSK4 was also reduced in the HMDMs compared to iMacs (Fig. 4F). NLRP1 interacts with ASC to activate caspase-1 while CARD8 interacts directly with caspase-1(48), consequently ASC specks should only be present if NLRP1 is activated. We therefore imaged ASC specks in iMacs treated with Talabostat. ASC speck formation is observed in all cells treated with Talabostat with the largest increase in specks observed in unprimed and Pam3CSK4-primed conditions (Fig. 4 G, H) similar to the LDH response. Therefore, in iMacs stimulation with Talabostat appears to activate the NLRP1 inflammasome.

**Fig. 4.**
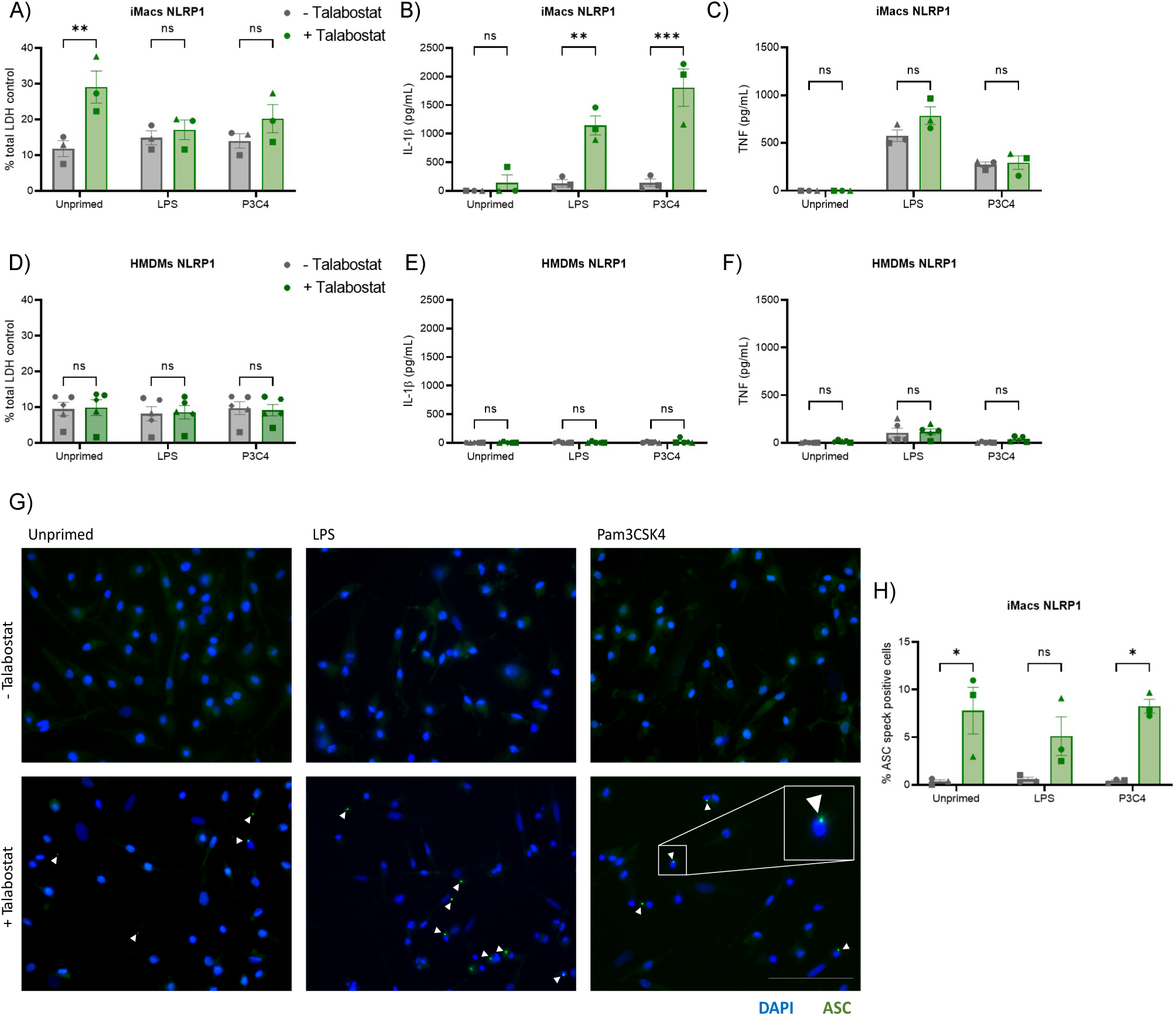
iMacs activate the NLRP1 inflammasome in response to Talabostat, unlike HMDMs. Quantification of **A)** LDH, **B)** IL-1β, and **C)** TNF release from iMacs in unprimed cells and following stimulation with 100 ng/ml LPS or 1 μg/ml Pam3CSK4 for 4 h with Talabostat (10 μM) treatment for 24 hours. Quantification of **D)** LDH, **E)** IL-1β, and **F** TNF release from HMDMs in unprimed cells and following stimulation with 100 ng/ml LPS or 1 μg/ml Pam3CSK4 for 4 h with Talabostat (10 μM) treatment for 24 hours. **G)** Representative images from iMacs in unprimed cells and following stimulation with 100 ng/ml LPS or 1 μg/ml Pam3CSK4 for 4 h with 10 μM Talabostat for 24 h stained with DAPI (blue) and ASC (green) with ASC specks highlighted with white arrows. Scale bar = 100 μM. **H)** Quantification of ASC speck positive cells. Data are represented as mean +/-SEM, n = 3 independent replicates, *p < 0.05, **p < 0.01, ***p < 0.001 by Two-way ANOVA with Šídák’s multiple comparisons test.

## Discussion

While some previous studies had observed inflammasome activation in iMacs, our aim here was to comprehensively characterize a range of inflammasome responses in iMacs and to benchmark them with HMDMs as the current gold standard model for human inflammasome signaling. In terms of the expression of macrophage markers and levels of phagocytosis, the iMacs generated in this study were very comparable to HMDMs. Previous work has characterized transcriptional responses to LPS(24) in iMacs, but as our interest is in protein complexes we used a proteomics approach. We found that iMacs generally express high levels of inflammasome sensors and other inflammatory proteins and thus have the capacity to form multiple inflammasomes. Our iMac proteomics dataset will be of significant interest for others interested in examining innate immune pathways using this cell model.

We found that NLRP3 was efficiently activated in iMacs, however, ATP induced a significantly lower inflammasome response compared to nigericin both in the iMacs and HMDMs. Extracellular ATP is an alarmin released from necrotic cells and functions as a DAMP to trigger NLRP3 activation(42). This lower response to ATP, could be influenced by the fact that the ATP receptor, P2X7R, is highly polymorphic and the KOLF2-C1 cells may have relatively low levels of P2X7R expression(49). In addition, Bezbradica et al., (2016) previously reported that sterile stimuli like ATP induce a significantly lower level of NLRP3 activation relative to microbial stimuli in HMDMs(42). The enhanced inflammasome response to microbial stimuli likely reflects the physiological requirement for faster and stronger immune responses to infection to limit pathogen dissemination. Indeed, we observed that iMacs activate NLRP3 in response to nigericin without any priming stimulus. Priming-independent NLRP3 activation by nigericin has previously been reported in human monocytes(50) but not in macrophages. HMDMs do not respond to nigericin alone so this is a distinct response in iMacs, potentially reflecting their more tissue resident-like ontogeny. Further work is needed to confirm whether this lower threshold for NLRP3 activation is consistent between iMacs and primary human tissue resident macrophages. Interestingly, TLR ligands alone can also activate NLRP3 in human monocytes(51), but we do not observe NLRP3 activation in response to LPS or Pam3CSK4 in iMacs.

Our iMacs also demonstrated a robust response to both NCI and NAIP/NLRC4 activation similar to HMDMs. This included high levels of pyroptosis in both primed and unprimed cells in response to both LPS transfection and Lfn-BsaL stimulation, confirming that priming is not necessary for NCI or NAIP/NLRC4 activation in human macrophages. Priming is required in mouse macrophages as caspase-11 is not constitutively expressed and must be upregulated(52). In human monocytes caspase-4 has been found to be constitutively expressed(53). Some studies have however reported a requirement for IFN-γ priming in LPS transfection-induced NCI activation in human epithelial cells and HMDMs to induce expression of granulocyte-binding proteins (GBPs) that are necessary for caspase-4 activation in response to cytosolic LPS(54). In iMacs, caspase-4 is constitutively expressed which may partly explain the dispensable requirement for priming. Mass spectrometry detected GBP1-4 expression in unstimulated iMacs, with an increase in GBP1 and GBP2 expression following 6 and 24 hour LPS stimulation. Therefore, constitutive GBP expression appears to be sufficient for NCI activation in iMacs.

We observed significant differences in the iMac response to NLRP1/CARD8 activation relative to HMDMs. In iMacs Talabostat causes pyroptosis and IL-1β release while HMDMs are unresponsive as previously reported(47). We observed that ASC specks were formed in Talabostat treated iMacs suggesting this was due to NLRP1 activation as CARD8 does not form ASC specks(48). Mouse NLRP1 is mostly expressed in myeloid cells such as macrophages, but human NLRP1 is predominantly found in keratinocytes and epithelial cells while CARD8 is more common in leukocytes(10,47). Our mass spectrometry analysis identified expression of both NLRP1 and CARD8 in the iMacs suggesting they express these functional inflammasomes. Therefore, iMacs can be an attractive model for studying NLRP1 and CARD8 compared to HMDMs. Future experiments can further explore the NLRP1 response using NLRP1-specific stimuli, such as viral dsRNA and viral infection and CARD8-specific stimuli, such as the aminopeptidase inhibitor, CQ80(38,55).

The “two-step” activation dogma of inflammasome activation involving priming and activation was developed based on mouse macrophage studies. Our results demonstrate that there is no requirement for priming of the NLRP3, NAIP/NLRC4, NLRP1, or non-canonical inflammasomes in human iMacs. Priming by TLR ligands simply increases pro-IL-1β which is necessary for measuring IL-1β release as a read-out of inflammasome activation. This increased sensitivity to inflammasome stimuli likely reflects a necessity to provide rapid and robust anti-microbial responses in tissues. In mice inflammasome activation is restricted by the necessity for two-step activation while humans may have evolved additional regulatory mechanisms such as COP and POP proteins to restrain excessive and potentially damaging inflammasome signaling.

In summary, we have shown that iMacs are an excellent model to study a range of inflammasomes. Unlike HMDMs, iMacs can be genetically modified opening up a wide range of new opportunities to study inflammasome function in macrophages including in disease settings. iMacs may also be a useful model for studying tissue resident macrophages, and protocols have been developed to differentiate iPSCs into specialized macrophages, such as microglia(56). Future work in terminally differentiated macrophages could therefore provide additional insight into the role of inflammasomes in specific tissues and disease states.

## Limitations

Experiments were performed using a single iPSC line, therefore the genetic background of the individual that the cells were originally derived from could influence the results. KOLF2-C1 cells have somatic mutations in genes such as AT-rich Interactive Domain 2 (ARID2)(39). ARID2 has roles in chromatin remodeling(57), but the potential effects of this mutation on inflammasome signaling are unknown. We also focused on testing responses to four well characterized and therapeutically relevant inflammasomes but did not examine other inflammasomes such as Pyrin or CARD8.

## Acknowledgements

This work was supported by funding from the Academy of Medical Sciences (SBR005\1104) to RCC; The Royal Society (RGS\R1\201127) to RCC; Biotechnology and Biological Sciences Research Council (BB/V016741/1) to RCC and BCC; and Northern Ireland Department for the Economy PhD studentships (to CMM, TJM, and ECM). We would like to thank Dr Subhankar Mukhopadhyay (King’s College London) for helpful discussions about iPSC-derived macrophage culture.

## Author contributions

Conceptualization: RCC, CMM. Funding acquisition: RCC, BCC. Investigation: CMM, MC, ECM, MA, TJM. Formal analysis: CMM, ECM, MA, BCC. Project administration: RCC. Supervision: RCC, BCC. Visualization: CMM, ECM. Writing – original draft: RCC, CMM. Writing – review & editing: CMM, MC, ECM, MA, TJM, BCC, RCC.

## Declarations of interest

R.C.C. is a co-inventor on patents and patent applications for NLRP3 inhibitors which have been licensed to Inflazome Ltd, Ireland. R.C.C. is a consultant for BioAge Labs, USA (since 2020), and serves on the Scientific Advisory Board of Viva in Vitro Diagnostics, Spain (since 2024). B.C.C. receives or has received research funding from Bruker Daltonics and Almac Discovery. All other authors have no competing interests.

## Supplemental information

Supplementary Table 1 contains protein group quantities from unstimulated, and LPS stimulated (6 h and 24 h) iMacs.

## Figure legends

**Supp. Fig 1.**
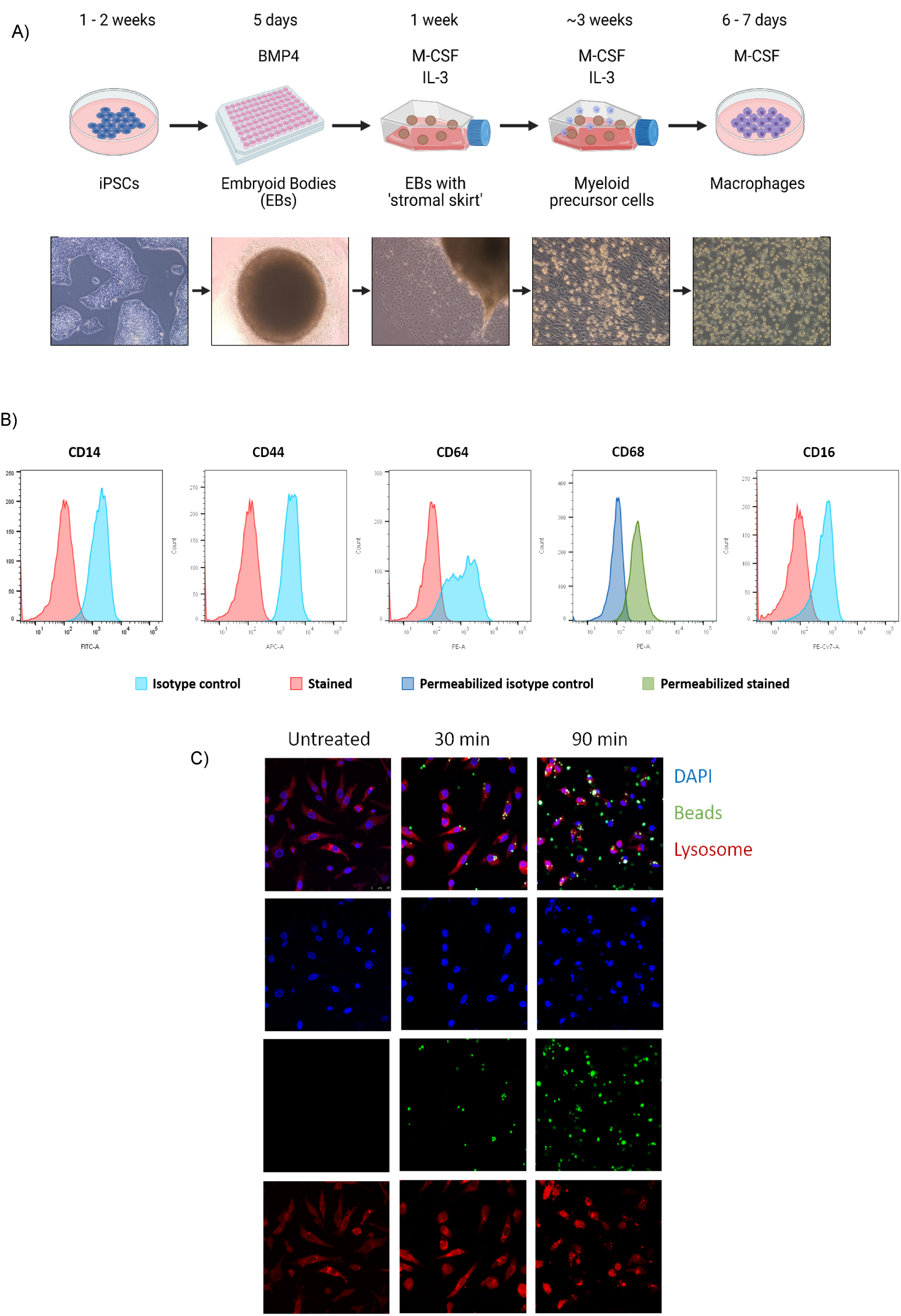
HMDMs express macrophage markers and are highly phagocytic. **A)** Schematic diagram illustrating the iMac differentiation protocol. **B)** Histograms showing representative expression of macrophage markers CD14, CD44, CD64, CD68, and CD16 in unstimulated HMDMs compared to relative isotype controls. **C)** 63 X (oil lens) Stellaris-5 confocal microscope representative images from HMDMs in untreated cells and following incubation with Latex amine modified polystyrene fluorescent beads (green) for 30 minutes or 90 minutes stained with DAPI (blue) and Lysotracker (red) with phagocytosed beads highlighted with white boxes. Scale bar = 100 μM.

**Supp. Fig 2.**
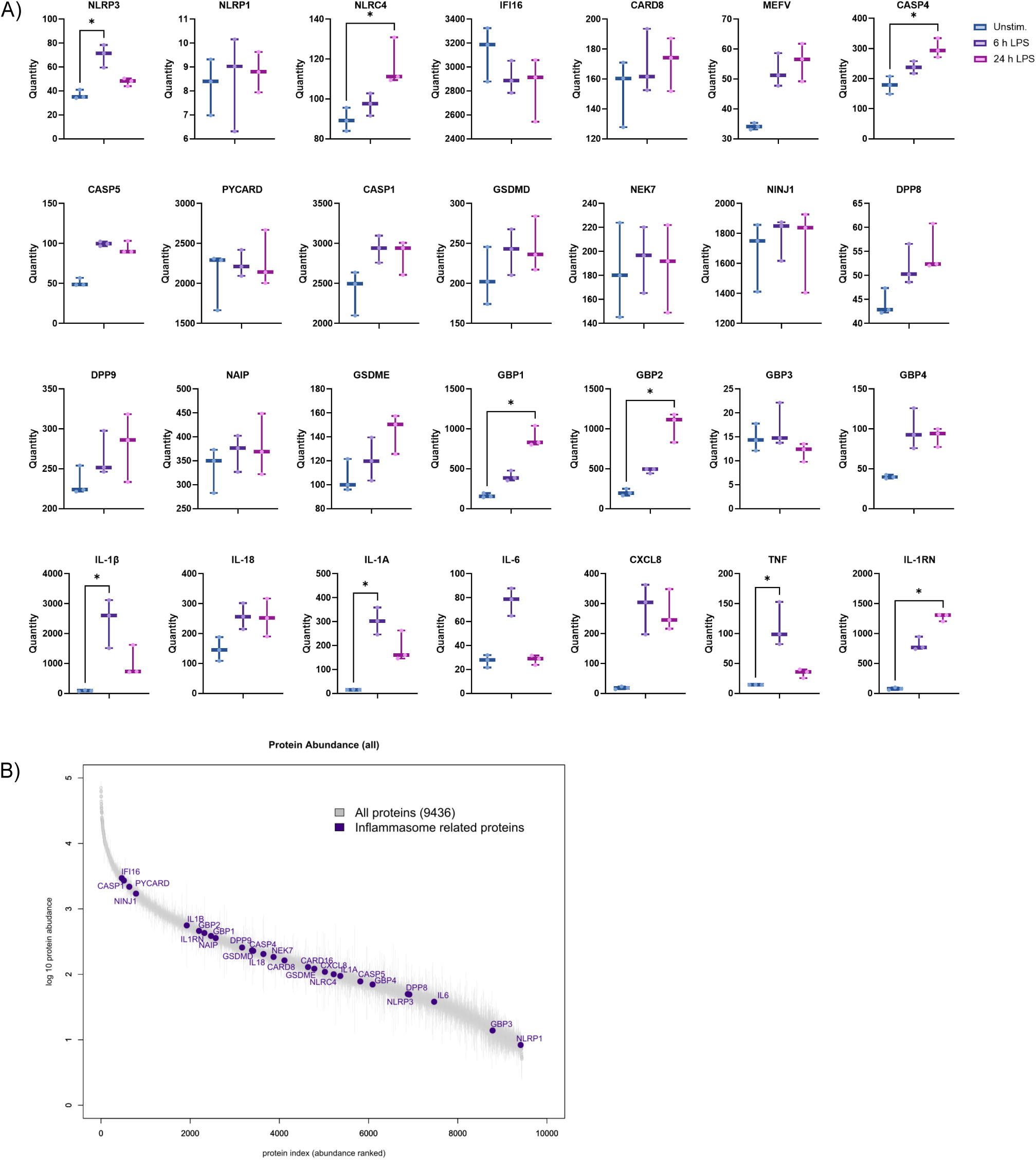
iMacs express a wide range of inflammasome- and inflammation-related proteins. **A)** Expression of inflammasome- and inflammation-related proteins in unstimulated iMacs or following stimulation with 100 ng/ml LPS for 6 h or 24 h. Data are represented as mean +/-SEM, n = 3 independent replicates, *p < 0.05 by Two-way ANOVA with Šídák’s multiple comparisons test. **B)** Plot depicting the ranked average Log10 abundance of proteins detected in iMacs across all experimental conditions and replicates (18 runs). Inflammasome related proteins are indicated.

**Supp. Fig 3.**
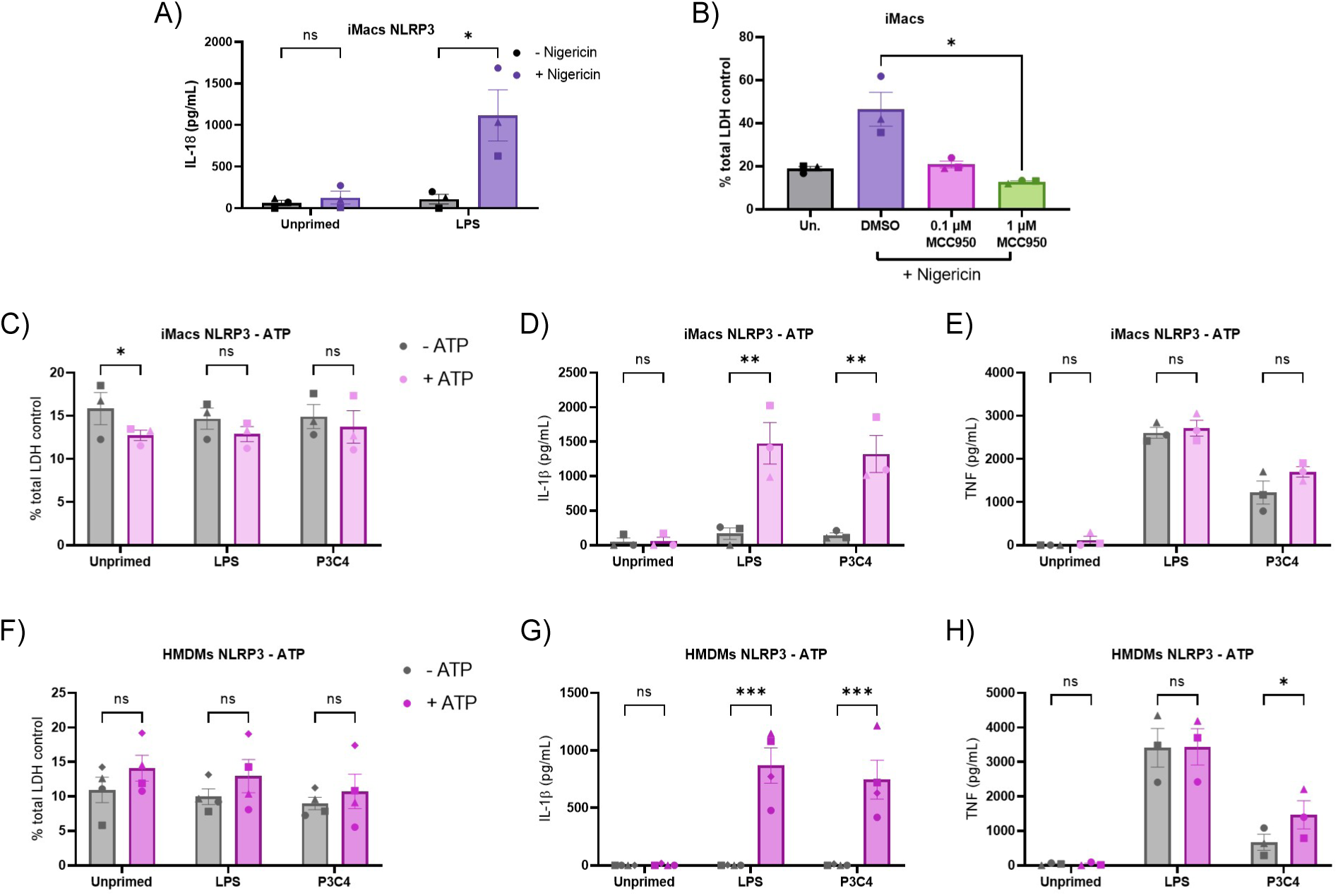
NLRP3 activation in iMacs and HMDMs. **A)** Quantification of IL-18 release from iMacs in unprimed cells and following stimulation with 100 ng/ml LPS for 4 h with 5 µM nigericin for 2 h. **B)** Quantification of LDH release from iMacs in untreated cells and cells pre-treated with MCC950 (0.1 μM or 1 μM) for 30 min before stimulation with 5 µM nigericin for 2 h. Quantification of **C)** LDH release, **D)** IL-1β, and **E)** TNF release from iMacs in unprimed cells and following treatment with 100 ng/ml LPS or 1 μg/ml Pam3CSK4 for 4 h with 5 mM ATP for 2 h. Quantification of **F)** LDH release, **G)** IL-1β, and **H)** TNF release from HMDMs in unprimed cells and following treatment with 100 ng/ml LPS or 1 μg/ml Pam3CSK4 for 4 h with 5 mM ATP for 2 h. Data are represented as mean +/-SEM, n = 3-4 independent replicates, *p < 0.05, **p < 0.01, ***p < 0.001, ****p < 0.0001 by Two-way ANOVA with Šídák’s multiple comparisons test.

**Supp. Fig 4.**
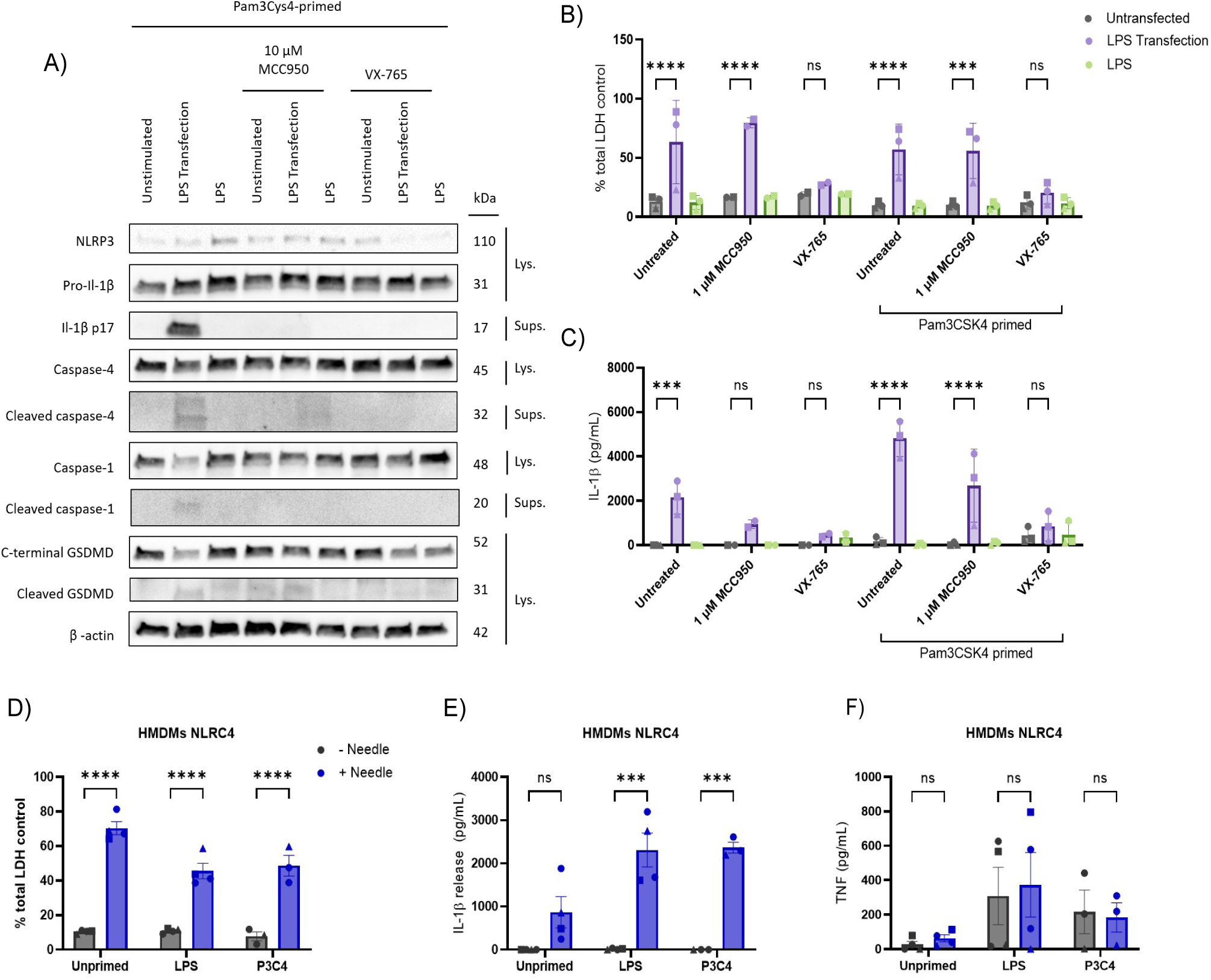
NCI and NLRC4 inflammasome activation in HMDMs. Quantification of **A)** LDH and **B)** IL-1β release from HMDMs in unprimed and Pam3CSK4 primed cells (1 μg/ml, 16 hours) and **C)** Western blots in from primed HMDMs following LPS transfection or LPS stimulation for 5 hours and treated with MCC950 (1 μM or 10 μM) or VX-765 (20 μM) prior to transfection. Quantification of **D)** LDH, **E)** IL-1β, and **F)** TNF release from iMacs in unprimed cells and following stimulation with 100 ng/ml LPS or 1 μg/ml Pam3CSK4 for 4 h with Lfn-BsaL (1 ng/ml) plus anthrax protective antigen (PA, 250 ng/ml) for 2 hours. Data are represented as mean +/-SEM, n = 3-4 independent replicates, *p < 0.05, ***p < 0.001, ****p < 0.0001 by repeated measures ANOVA with Tukey’s multiple comparisons test.

## STAR Methods

### Culture and differentiation of induced pluripotent stem cell-derived macrophages

KOLF2-C1 iPSCs were obtained from the Wellcome Sanger Institute (Hinxton, UK). KOLF2-C1 iPSCs were thawed and plated onto vitronectin (ThermoFisher, Cat #A14700) coated plates in StemFlex (ThermoFisher, Cat #A3349401) with 10 μM Rho-kinase (Rock) inhibitor Y-27632 Dihydrochloride (Merck, Cat #Y0503). Cells were cultured in tissue culture incubators at 37°C with 5% CO2 in a humidified atmosphere. Medium was changed daily (without Rock inhibitor) until cells were 70-80% confluent. Cells were passaged using 0.5 mM PBS-EDTA and plated onto a vitronectin-coated 10 cm2 tissue culture dishes in StemFlex medium with 10 μM Rock inhibitor.

To generate embryoid bodies (EBs), iPSCs were harvested and seeded onto a low-adherent, U-bottom 96 well plate at 1x10^5^ per well in 100 μl in StemFlex medium supplemented with 50 ng/mL BMP-4 (R&D Systems, Cat #314-BP010) and 10 μM Rock inhibitor. The 96 well plates were centrifuged for 3 minutes at 800 G, 4°C and incubated for 5 days to form EBs.

The EBs were transferred to gelatine (Sigma-Aldrich, Cat #G1890) coated tissue culture flasks in myeloid precursor base medium (X-VIVO 15 Serum Free Medium; Lonza, Cat #BE02-060F, supplemented with 2 mM GlutaMAX; ThermoFisher, Cat #35050061 and 100 IU/ml penicillin-streptomycin; ThermoFisher, Cat #15140122). Media was changed every 4 – 5 days. Approximately 3 – 4 weeks after transferring the EBs to the gelatine-coated plates, they began to produce myeloid precursor cells which were then harvested and plated in iMac medium (RMPI 1640; ThermoFisher, Cat #21870076 supplemented with 1-% fetal bovine serum; ThermoFisher, Cat #26140079, 2 mM GlutaMAX, 100 IU/ml penicillin-streptomycin, and 100 ng/mL M-CSF; ImmunoTools, Cat #11343118) and differentiated for 7 days. Differentiated iMacs were harvested using Lidocaine monohydrate (Sigma-Aldrich, Cat #L5647) and plated onto 96-, 24-, and 6-well plates at 0.5x10^5^ in 100 μl, 2x10^5^ in 500 μl, and 1x10^6^ cells per well respectively in iMac medium.

### Inflammasome activation

To activate NLRP3, cells were primed with 100 ng/ml ultrapure LPS from E. coli K12 (InvivoGen, Cat #tlrl-peklps) or 1 μg/ml synthetic triacylated lipopeptide Pam3CSK4 (InvivoGen, Cat #tlrl-pms) for 4 hours and treated with 5 μM Nigericin sodium salt (AdipoGen, Cat #AG-CN2-0020-M005) or 5 mM Adenosine 5ʹ-Triphosphate (ATP) disodium salt (Merck Life Science, Cat #1191-1GM) for 2 hours. To activate NLRP1, cells were primed and treated with 10 μM Talabostat mesylate (R & D Systems, Cat# 3719/10) for 24 hours. To activate NAIP/NLRC4, cells were primed and treated with Lfn-BsaL (1 ng/ml, a fusion protein of the N-terminal domain of the Bacillus anthracis lethal factor and BsaL needle protein from Burkholderia pseudomallei, InvivoGen, Cat #tlrl-ndl) plus 250 ng/ml Anthrax Protective Antigen (PA), Recombinant from Bacillus anthracis (Quadratech Diagnostics, Cat #LL-171E) for 2 hours. To activate the non-canonical inflammasome, iMacs were primed overnight with 1 μg/ml Pam3CSK4 (around 16 hours) then fresh media was added and LPS was transfected using Lipofectamine LTX Reagent with PLUS Reagent (ThermoFisher, Cat #A12621) and centrifuged at 1000 G for 10 min at room temperature before being incubated for 5 hours. To inhibit NLRP3, cells were pre-treated with 1 μM or 10 μM MCC950 (Caltag Medsystems, Cat #AG-CR1-3615-M005) for 15 minutes prior to treatment with nigericin or transfection with LPS. To inhibit caspase-1, cells were pre-treated with 20 μM VX-765 (MedChemExpress, Cat #HY-13205 -5 mg) for 15 minutes.

### Flow cytometry

iMacs were resuspended and 0.5x106 added per flow tube in 1 ml DPBS. All cells were stained with eBioscience Fixable Viability Dye eFluor 506 (ThermoFisher, Cat #65-0866-14) at a dilution of 1:500 for 10 minutes at room temperature then washed twice in DPBS. Blocking solution (Human TruStain FXX, Biolegend, Cat #422302) was added at a dilution of 1:20 for 30 minutes in the dark at room temperature and cells were washed twice in DPBS. For intracellular antibodies, cells were resuspended in 200 μl fixation buffer (eBioscience™ Intracellular Fixation & Permeabilization Buffer Set, ThermoFisher, Cat #88-8824-00) for 30 minutes at room temperature. The cells were centrifuged for 3 minutes at 800 G and resuspended in 200 μl permeabilization buffer (1:10 from the 10X stock, diluted in ddH20, eBioscience™ Intracellular Fixation & Permeabilization Buffer Set, ThermoFisher, Cat #88-8824-00). The cells were centrifuged for 5 minutes at 500 G and the cell pellet resuspended in 500 μl permeabilization buffer. The appropriate antibody/isotype control concentration was added, and the cells were incubated for 30 minutes in the dark at 4°C. For extracellular antibodies, 500 μl DPBS was added to each tube with the appropriate antibody/isotype concentration and incubated for 30 minutes in the dark at 4°C. Cells were stained with antibodies and isotype controls listed in the key resources table. The flow tubes were centrifuged for 5 minutes at 500 G, room temperature and washed twice with DPBS. Cells stained for extracellular antibodies were resuspended in DPBS and cells stained for intracellular antibodies were resuspended in permeabilization buffer. Samples were run on the FACS Canto II cytometer (Becton Dickenson, UK) and results analyzed using FlowJo software (BD Biosciences, UK).

### Phagocytosis Assays

iMacs were plated at 1.25x105 per well in 24 well plates on coverslips in iMac medium and incubated overnight. RPMI was removed and replaced with OptiMEM (ThermoFisher, Cat #31985070). Latex amine-modified polystyrene, fluorescent yellow-green beads (Sigma-Aldrich. Cat #L1030) were diluted 1:10 in DPBS and added 1:50 per well. iMacs were incubated with the fluorescent beads for 30 or 90 minutes. OptiMEM was removed and cells were washed in DPBS. Fresh OptiMEM was added and 0.5 μM LysoTracker Red DND-99 (ThermoFisher, Cat #L7528,) added for 30 minutes.

Upon completion of the experiment, cells were washed in DPBS and fixed in 4% paraformaldehyde (PFA) for 15 minutes. PFA was aspirated and coverslips were stored in PBS at 4°C until staining.

Coverslips were washed 3 times in PBS then removed from the 24 well plates and kept in a humidified chamber. Coverslips were permeabilized with 0.1% Triton X-100, washed in PBS and then PFA was quenched with 50 mM ammonium chloride. Coverslips were washed then blocked with 0.5% BSA/PBS for 30 minutes before mounting in mounting media containing DAPI on glass microscope slides. The mounted coverslips were sealed with clear nail polish and stored at 4°C until imaging. Coverslips were imaged at 63 X (oil lens) using the Stellaris-5 confocal microscope. An overview of each coverslip was taken using the automated spiral scan on the Leica LAS X software. Images were taken at a 16-bit resolution using a 1024x1024 scan format. Representative images were analyzed using Fiji. The percentage of DAPI+bead+ cells was calculated.

### Mass spectrometry

iMacs were plated at 1x106 in 6 well plates and challenged with LPS (100 ng/ml) or Pam3CSK4 (1 μg/ml) for 24 hours. The cells collected in ice-cold PBS into 1.5 ml tubes and centrifuged at 13,000 RPM at 4°C for 10 minutes. The cell pellets were frozen at -80°C. The cells were lysed with 1% SDC lysis buffer (1% SDC, 10 mM TCEP, 40 mM 2-chloroacetamide (CAA), 100 mM Tris pH 8.5) and heated for 10 minutes at 95°C, with shaking (800 RPM). Samples were sonicated for 2 minutes then centrifuged at maximum speed for 15 minutes and the supernatants stored at −20°C. Preparation for mass spectrometry was carried out using a protein aggregation capture (PAC) protocol adapted from (58) using the Opentrons OT-2 liquid handling system. Lysates were added to MagReSyn Hydroxyl beads (ReSyn Biosciences) for aggregation for 20 minutes, with a mixing step at 10 minutes. The bead–protein aggregates were washed twice with Acetonitrile (ACN) and once with 50 mM triethylammonium bicarbonate buffer (TEAB) pH 8.5 at a slow speed. Proteins were digested with 50 mM TEAB containing Lys-C and Trypsin at an enzyme-to-substrate ratio of 1:100 and 1:50, respectively for 4 h at room temperature. Evotips (Evosep) were activated by soaking in isopropanol for 10 seconds, then pre-conditioned by pushing through 20 µL of 0.1% FA. 50% of the peptides were loaded onto Evotips and mass spectrometry was performed on a timsTOF HT (Bruker) quadrupole time-of-flight mass spectrometer integrating trapped Ion Mobility (IM) separations coupled online via a Captivespray electrospray source (Bruker) to Evosep One liquid chromatography system(59,60). Peptides were separated on an integrated packed emitter column (15 cm × 75 μm ID – Coann) packed with 1.5 μm Reprosil Saphir beads (Dr. Maisch) using whisper 40 SPD method. Data was acquired using a diaPASEF method consisting on 12 cycles including a total of 24 windows, covering m/z from 300 to 1400 Da with two mobility windows each, covering the ion mobility range (1/K0) from 0.70 to 1.42 V s/cm2. The CaptiveSpray source was kept at a capillary voltage of 1,600 V, with a dry gas flow rate of 3.0 L/min at 180 °C. Accumulation and ramp times were set to 100 ms, resulting in 180 ms per MS¹ or MS² (DIA-PASEF) scan, excluding transfer time. The collision energy was programmed as a function of ion mobility, following a straight line from 20 eV for 1/K0 of 0.6 V s/cm2 to 59 eV for 1/K0 of 1.6 V s/cm2. To calibrate TIMS, the elution voltage was set to linear mode to obtain 1/K0 ratios using three ions (m/z of 622, 922, 1222) from the ESI-L Tuning Mix (Agilent). An additional gas phase fractionation run (GPF) of a pooled sample was included to boost library coverage(61).

Mass spectrometry data was analyzed in Spectronaut version 19.1 (Biognosys) using the using a direct DIA workflow with default BGS settings against UniProtKB human sequence database UP000005640 (downloaded January 2024 - 20,413 entries); specific Trypsin/P digestion, 2 missed cleavages, peptide length 7–52, carbamidomethyl cysteine set as fixed modification, oxidation of methionine and N-terminal methionine excision as variable modifications allowing up to five variable sites per peptide. For PSMs, peptides, and proteins, the false discovery rate was controlled at 1%. Precursor filtering required a peptide–spectrum match q-value of ≤ 0.01 and a minimum posterior error probability of 0.2. Protein identifications met q-value thresholds of 0.01 for experiment-wide assessment and 0.05 for run-wide assessment. Protein grouping was performed using the IDPicker algorithm. Protein quantification was performed using the MaxLFQ algorithm on peak areas, followed by cross-run normalization, without imputation of missing values. UniProtKB human sequence database UP000005640 (83,385 entries). The protein quantities table was exported using a pivot report in Spectronaut. Downstream plots were generated in R, using base R and the following packages: ggplot2, ggrepel, pheatmap.

### Lactate Dehydrogenase (LDH) assay

LDH assay (Cytotoxicity Detection Kit Roche, Sigma Aldrich, Cat #11644793001) was performed following manufacturer’s instructions to assess cell death.

### Enzyme-Linked Immunosorbent Assay (ELISA)

Human IL-1 beta/IL-1F2 DuoSet ELISA (R&D Systems, Cat #DY201), Human TNF-alpha DuoSet ELISA (R&D Systems, Cat #DY210) and human IL-18 ELISA (ThermoFisher, Cat #BMS267-2MST) were performed following manufacturer’s instructions.

### Western blot

Cell lysates were harvested in 50 μl cell lysis buffer (66 mM Tris HCl pH 7.4% SDS) with 1:10,000 BaseMuncher (Expedeon Cat #BM0025). 1X NuPAGE LDS sample buffer (Invitrogen, Cat #NP0007) plus Dithiothreitol (DTT, Merck Life Science, Cat #20-265) was added to each sample. Samples were boiled for 5 minutes at 95°C then run on 4–20% Mini-PROTEAN® TGX™ Precast Protein Gels (BioRad, 10 wells, Cat #4561093, 12-wells, Cat #456109, or 15-wells, Cat #4561096) and were separated for 60 – 80 minutes at 120 V alongside Precision Plus Protein Kaleidoscope Standard (BioRad, Cat #1610375EDU). Proteins were transferred to a nitrocellulose membrane using the Trans-Blot® Turbo™ Transfer System (BioRad) at 25V for 7 minutes. Membranes were then blocked in 5% dried milk powder in TBS-Tween at room temperature for 1 hour. Membranes were then incubated with one of the following antibodies: caspase-1 (Cell Signaling Technology, Cat #3866), cleaved caspase-1 (Cell Signaling Technology, Cat #4199S), caspase-4 (Cell Signaling, Cat #4450S), cleaved GSDMD (Abcam, Cat #ab215203), C-terminal GSDMD (Abcam, Cat #ab210070), NLRP3 (Cell Signaling, Cat #15101), IL-1β (R&D Systems, Cat #AF401-NA), or β-actin (Sigma, Cat #A2228) overnight on a rocker 4°C. After incubation, the membranes were washed and incubated with the appropriate secondary antibody: anti-rabbit (Jackson Immunoresearch, Cat #111-035-003), anti-mouse (Jackson Immunoresearch, Cat #515-035-003), or anti-goat (Jackson Immunoresearch, Cat #705-035-147). Proteins were detected by incubating membranes with Clarity ECL (Biorad, Cat #170-5061) and the fluorescence imaged using the GeneSys imager.

### ASC speck staining and imaging

Cells were plated at 1x10^5^ per well in their culture medium overnight in 24 well plates with coverslips added. Medium was then replaced with OptiMEM per well. The indicated stimuli were added for times listed then cells were washed in PBS and fixed in 4% paraformaldehyde (PFA) for 15 minutes. PFA was aspirated and coverslips were stored in PBS at 4°C until staining.

Coverslips were washed 3 times in PBS then removed from the 24 well plates and kept in a humidified chamber. Coverslips were permeabilized with 0.1% Triton X-100, washed in PBS and then PFA was quenched with 50 mM ammonium chloride. Coverslips were washed then blocked with 0.5% BSA/PBS for 30 minutes. Primary anti-ASC antibody (BioLegend, Cat #676502) was diluted 1:50 in 0.5% BSA/PBS and added for 60 minutes in the dark then coverslips were washed in PBS 4 times for 5 minutes. Alexa Fluor™ 488 Donkey anti-mouse (Invitrogen, Cat #A21202) was diluted 1:800 in 0.5% BSA/PBS and added for 60 minutes in the dark. The coverslips were washed in PBS 4 times for 5 minutes before mounting in mounting media containing DAPI on glass microscope slides. The mounted coverslips were sealed with clear nail polish and stored at 4°C until imaging. Coverslips were imaged using the Leica LAS X software and the DM5500 fluorescent microscope at 20X and 40X. ASC specks were quantified using Fiji and the percentage of DAPI+ASC+ positive cells was calculated.

### Statistics

For datasets comparing the means of three or more groups with one independent variable, statistical significance was checked through performing a One-Way ANOVA with Dunnett’s post-test. For datasets comparing the means of three or more groups with more than one independent variable, statistical significance was checked through performing a Two-Way ANOVA with Sidak’s or Tukey’s post-test. A power calculation was not performed. Graphs were generated using GraphPad Prism 9. Data are presented as mean +/-SEM, n number indicated in figure legends; n.s. not significant, *p < 0.05, **p < 0.01, ***p < 0.001, ****p < 0.0001. Specific statistical tests used are described in detail in figure legends

### Data availability

The mass spectrometry proteomics data have been deposited to the ProteomeXchange Consortium via the PRIDE (62) partner repository with the dataset identifier PXD068076. All other raw data are available upon request.

